# Mapping of single-cell landscape of acral melanoma and analysis of molecular regulatory network of tumor microenvironment

**DOI:** 10.1101/2021.08.26.457785

**Authors:** Zan He, Zijuan Xin, Xiangdong Fang, Hua Zhao

**Author notes:** Equal first author (Zan He, Zijuan Xin).

## Abstract

Melanoma is a type of skin malignant tumor with high invasiveness, high metastasis, and poor prognosis. The incidence of melanoma continues to increase. Among them, the subtype of acral melanoma (AM) is more common in Asian populations. AM has higher degree, low immunotherapy response rate. With the help of single-cell sequencing technology provides new technical means for tumor microenvironment research, so that we can more easily explore specific tumor types suitable immunotherapy targets. However, no complete single-cell level differentiation map exists for the AM tumor microenvironment (TME). In this study, we used AM related sample and used the 10× Genomics single-cell transcriptome platform to draw a specific single-cell map of AM, understand the cell composition of AM, and analyze the interaction and molecular regulation of AM TME. Nine cell types were identified, of which malignant cells accounted for the largest proportion, followed by fibroblasts. And the cell interaction network shows that malignant cells, macrophages, B, T and fibroblasts play an important role in AM TME. Our research provides systematic theoretical guidance for the diagnosis and treatment of acral melanoma.

## Introduction

Malignant melanoma is a malignant tumor derived from melanocytes. In recent years, the incidence of malignant melanoma continues to rise with an annual growth rate of 3%-5%. There are about 300,000 new cases and 60,000 deaths every year, posing a serious threat to people’s health (1). In patients with confirmed or suspected in-situ melanoma, surgical resection before metastasis has been found to be the first treatment option with a good prognosis. Metastatic melanoma, while accounting for only 1% of all cancer cases, is the deadliest (2). Patients are prone to relapse after surgery, resulting in a poor prognosis and a 5-year survival rate of less than 10% (3). Melanoma is usually divided into four types: cutaneous melanoma (mostly in the head, neck, trunk and limbs), acral melanoma (mostly in the palms, soles of feet and nail beds where there are no hair follicles), mucosal melanoma and uveal melanoma. Different from the epidemiological characteristics of less than 10% of AMs in western white population, the proportion of AMs in Asians is as high as 70% (4, 5), and AMs have been considered as a subtype of melanoma with poor prognosis and poor immune efficacy (6, 7). Compared with Caucasian patients, Chinese AM patients were more advanced, with larger tumor diameter and thicker tumor thickness, ulcer and high Clark grade. The histopathological features of AM are different from those of non-acral type. The Breslow thickness of AM is thicker, more frequent with ulcers, vertical growth, and lymph node metastasis.

The invasion and metastasis of malignant tumor are the pathologic basis of tumor recurrence, disease deterioration and death. Tumor invasion and metastasis is a continuous and progressive, multi-factor and multi-step process coordinating with TME. The immune system recognizes tumor antigens and kills tumor cells, but it is not enough to eliminate tumors that have formed in the body. Solid tumor is a complex tissue composed of not only tumor cells, but also stromal cells, inflammatory cells, vascular system and extracellular matrix (ECM), which are collectively defined as TME (8).

Single-cell sequencing is a study of the genome and transcriptome at the single-cell level (9). Through genome-wide or RNA amplification, high-throughput sequencing can reveal the gene structure and gene expression status of individual cells, reflecting inter-cell heterogeneity. Compared with traditional RNA-seq, single-cell sequencing(scRNA-seq) is more suitable for the analysis of TME components and heterogeneous populations. Its application in the study of TME has provided an unprecedented solution to the cellular and molecular complexity of it, thus deepening our understanding of heterogeneity, plasticity, and the complex cross-over interactions between different cell types in TME. With the continuous accumulation of scRNA-seq data sets, it will become an indispensable part of tumor immunology. It will continue to drive scientific innovation in precision immunotherapy and eventually be adopted in routine clinical practice in the foreseeable future. This new technique allows for better characterization of developmental lineages and differentiation states, which are critical to understanding the underlying mechanisms that drive the functional diversity of immune cells in TME.

We collected lesion biopsy samples from 5 clinical AM patients along with two adjacent paracancerous tissues and a metastatic lymph node sample, and performed 10×Genomics scRNA-seq and analysis respectively. Our study delineates a scRNA landscape of AM, and describes the molecular regulatory network of TME cells.

## Results

### Single-cell transcriptome sequencing and cell type identification

We collected 8 clinical tissuess from 5 patients with acral melanoma, including 5 samples of primary AM lesions (T), 2 samples of adjacent tissues(N), and 1 sample of lymph node metastasis tissue (L). After quality control and removal of batch effects, a total of 61,726 single cells were used for downstream analysis (Figure1.A, FigureS1, FigureS2.A-B). According to the CNV result by copyKAT (10), we initially identified aneuploid mutant cells as malignant cells, which are mainly derived from primary and lymph node metastasis tissues, and diploid mutant cells are identified as cells of other microenvironmental components (FigureS1.A-C). The microenvironment component cells were clustered into 15 clusters, and SingleR (11) was used to annotate them (FigureS1.D-F). All cells are defined as: malignant cells (MITF+, 21624 cells), cancer associated fibroblasts (CAFs) (COL1A1+ 17434 cells), T cells (CD3D+, 8322 cells), macrophages (C1QB+, 5520 cells), endothelial cells (PECAM1+, 4033 cells), neutrophils (S100A8+, 2519 cells), B cells (CD79A+, 1007 cells), epithelial cells (EPCAM+, 657 cells), keratinocytes (KRT14+, 607 cells) (Figure1.B, E). Then, according to the correlation of the sample clustering and the proportion of each type of cell in the sample, we found that sample clustering is related to the proportion of malignant cells (Figure1.F). At the same time, according to the progression of tumor, the proportion of stromal cells such as endothelial cells gradually decrease, the proportion of immune cells such as T cells gradually increase (Figure1.G).

**Figure 1.**
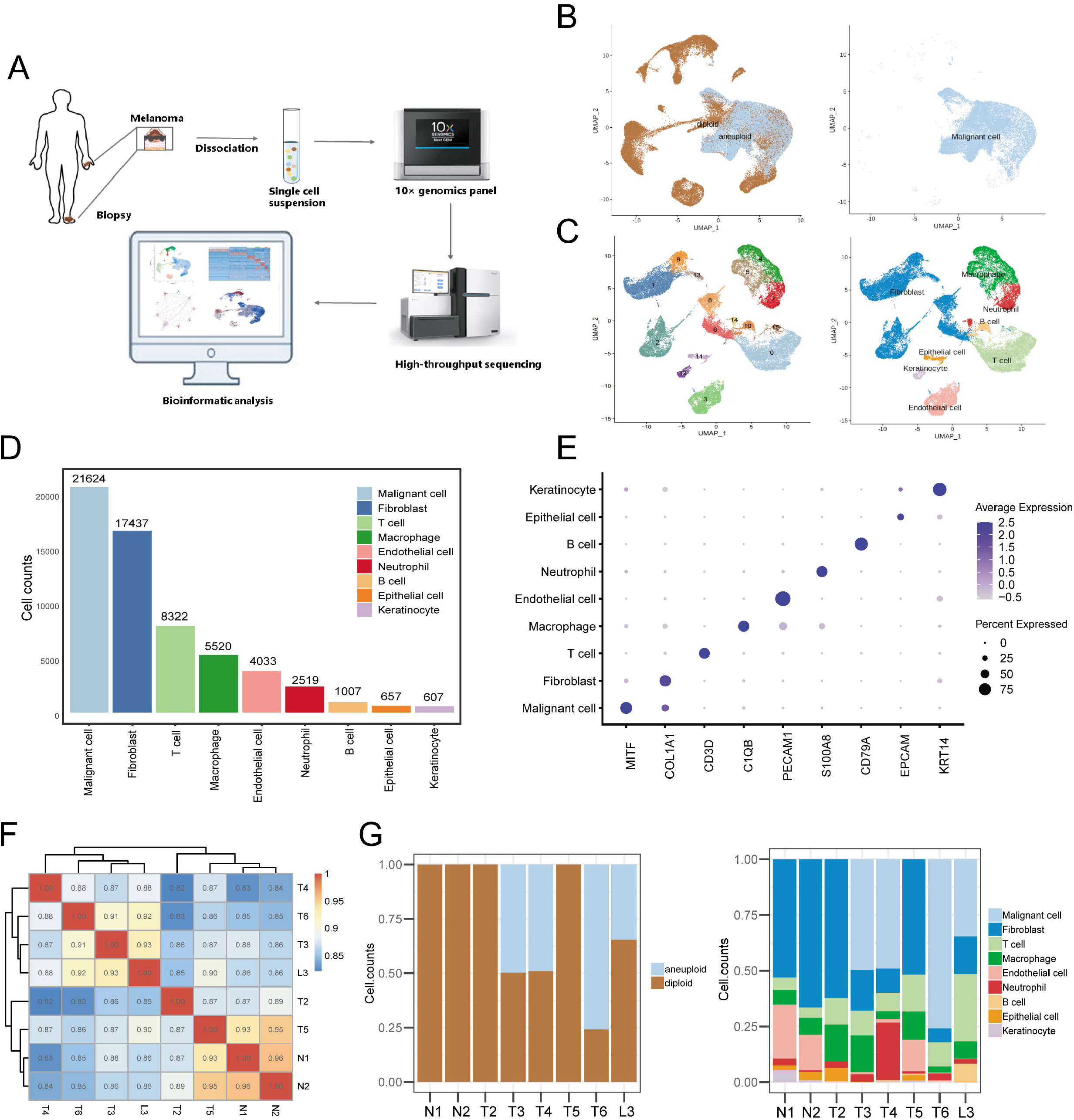
Cell composition in AM. A. Sample preparation, sequencing and bioinformatic analysis process. B. UMAP plot showing the Copykat results. C. UMAP plot of non-malignant cells labeled by cell cluster and cell type. D. cell counts bar plot of each cell type. E. Marker gene expression of each cell type, including dot size and color representing the characteristics of gene expression (pct.exp) and average scaling expression (avg. exp.scale) values. F. Sample pearson correlation coefficient heatmap. G. The distribution of the cell composition of each patient sample.

### TME Cell Interaction Network

According to the gene expression of the receptor-ligand pair, the cell interaction strength of primary tissues, adjacent tissues, lymph node metastasis tissue was inferred, and the cell interaction network was obtained by CellChat (12). The results showed that the communication between cells in the primary tissues was closer (Figure 2.A-C). In adjacent tissues, KIT and WNT cell interaction signal pathways were specifically identified, in which KIT signals were mainly secreted by endothelial and fibroblasts, and WNT signals were mainly secreted by keratinocytes. primary tissues specifically identified NT, ncWNT, IL1, and GDF cell interaction signaling pathways. NT, ncWNT, and GDF signals were mainly secreted by malignant cells, and IL1 signals were mainly secreted by neutrophils. Chemerin, NRG and GDF cell interaction signaling pathways were specifically identified in lymph node metastasis tissues. Chemerin and NRG signals were mainly secreted by malignant cells, and PSAP signals are mainly secreted by macrophages (Figure2.D, FigureS3.A-D). With the progression of tumor, the status of malignant cells in tumor associated macrophage(TAM) has gradually increased, and the interaction between malignant cells and macrophages, B cells, T cells and fibroblasts has gradually increased. These results suggest that malignant cells, macrophages, B cells, T cells and fibroblasts play more important roles in TME (Figure1.E, FigureS3.E).

**Figure 2.**
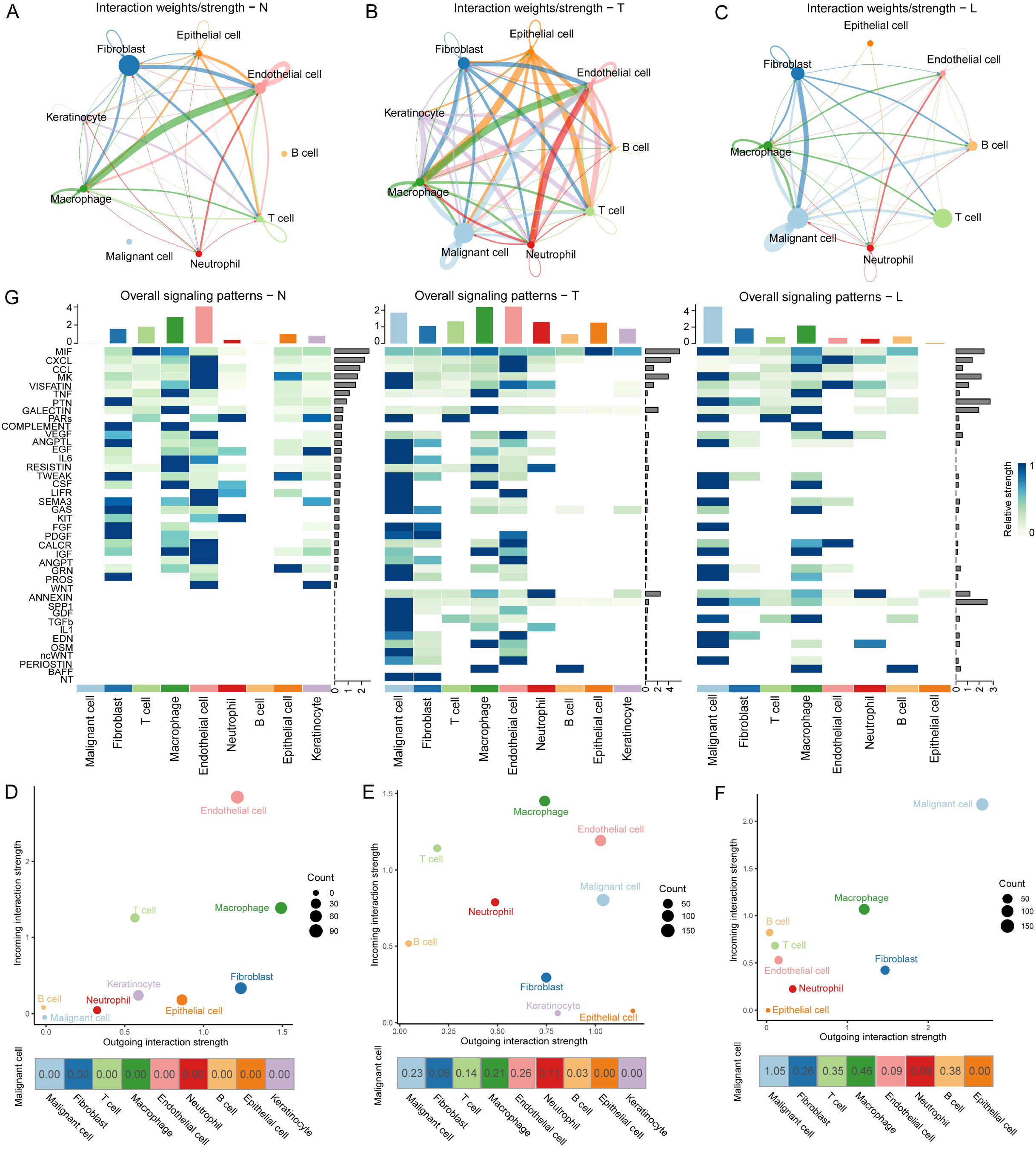
Cell-cell communicate analysis. A. Normal sample cell communication network. B. Tumor sample cell communication network. C. Lymph node metastasis sample cell communication network. D. Heatmap shows normal, tumor and lymph node metastasis cell interaction pathways identified by each cell type. E. The outgoing and incoming interaction strength of each cell type and Malignant cells and other types of cells in a normal samples. F. The outgoing and incoming interaction strength of each cell type and Malignant cells and other types of cells in tumor samples. G. The outgoing and incoming interaction strength of each cell type and Malignant cells and other types of cells in the lymph node metastasis sample.

### Pseudotime analysis of malignant cells

Malignent cell subpopulation analysis identified 6 cell subpopulations at different stages of differentiation. The pseudotime analysis inferred from splicing kinetics found that cells in cluster3 were in an earlier state, while cells in cluster1 were in a more terminal state (Figure3.A-B). Differential genes(DEGs) of cluster1 and 3 were enriched in GO function. We found that the highly expressed DEGs in cluster1 are related to behaviors such as extracellular matrix, response to TGFβ, regulation of epithelial cell proliferation, epithelial cell migration, etc. (Figure 3.C-D). We believed that the cells of cluster1 are in a highly malignant state. SCENIC can use random forest to identify the transcription factor (TF) co-expression network (13). It has identified the corresponding TF modules in cluster3 and cluster1 respectively. Among them, the TF TWIST1 and its corresponding co-expression module in the cluster with higher malignancy have the same AUC score and The expression level is higher (Figure3.E-F). Based on the above analysis, we suspect that TWIST1 may play an important regulatory role in the development of malignant cells by regulating the target genes in its co-expression module (Figure S4.G). Further GO function enrichment revealed that the genes in the TWIST1 co-expression module may be involved in the biological processes related to tumor deterioration such as epithelial mesenchymal transition (EMT) and response to TGFβ (Figure 3.H)

**Figure 3.**
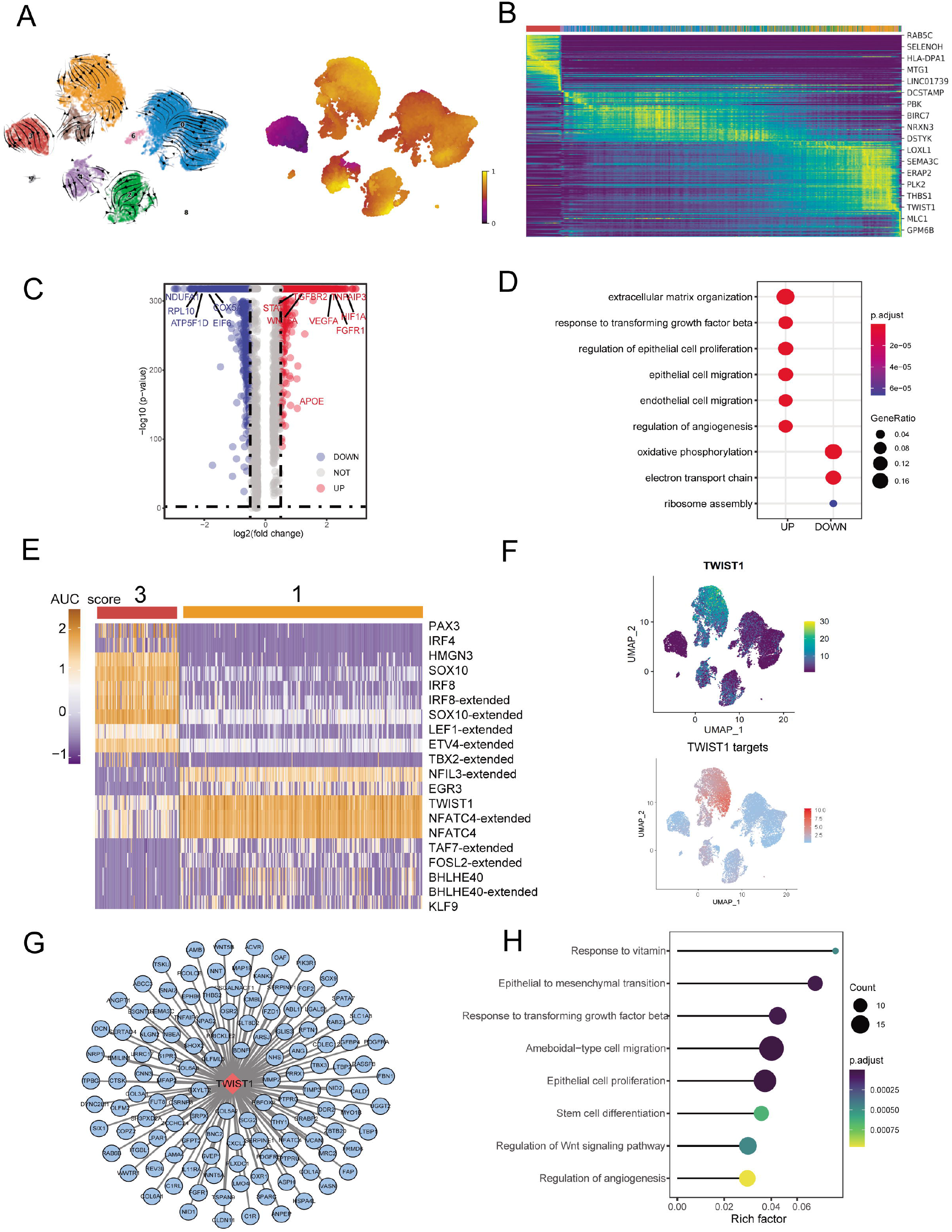
Malignant cell subpopulations analysis. A. UMAP plot shows the RNA velocities and the latent time of malignant cell subpopulations. B. Heatmap shows dynamic gene expression patterns accompanying the evolution of malignant cells. C. The volcano map shows the gene difference analysis between malignant subpopulations one and three. D. GO term enrich of DEG between malignant subpopulations one and three. E. Heatmap of the AUC scores of TF motifs estimated by SCENIC for each cell in subpopulations one and three. F. UMAP plot of each malignant cell TWIST1 and TWIST1 targets expression level. G. Network of TWIST1 and target genes. H. GO term enrichment results of TWIST1 targets.

### Two different subtypes of macrophages are identified in AM

The recluster analysis of macrophages distinguishes macrophages into two cell types, M1 (HLA-DQA2+) and M2 (CD163+), among which tumor-infiltrating macrophages are mainly M2 type (Figure 4.A). Combined with the key motifs identified by SCENIC, we realized that the activities of the three motifs IRF4 and SOX18 were down-regulated, and the activation of the STAT1, REL and NFKB1 motifs led to this M2 polarization process (Figure 4.B). Cell communication weight shows that there is a strong cell interaction between M2 cells and malignant cells, and there is also a certain cell interaction with M1 cells (Figure 4.C). Among them, M1 cells mainly secrete TNF, while M2 secretes TGFβ. In addition, M2 cells will also affect M1 through the Galectin signaling pathway (Figure 4.D-F).

**Figure 4.**
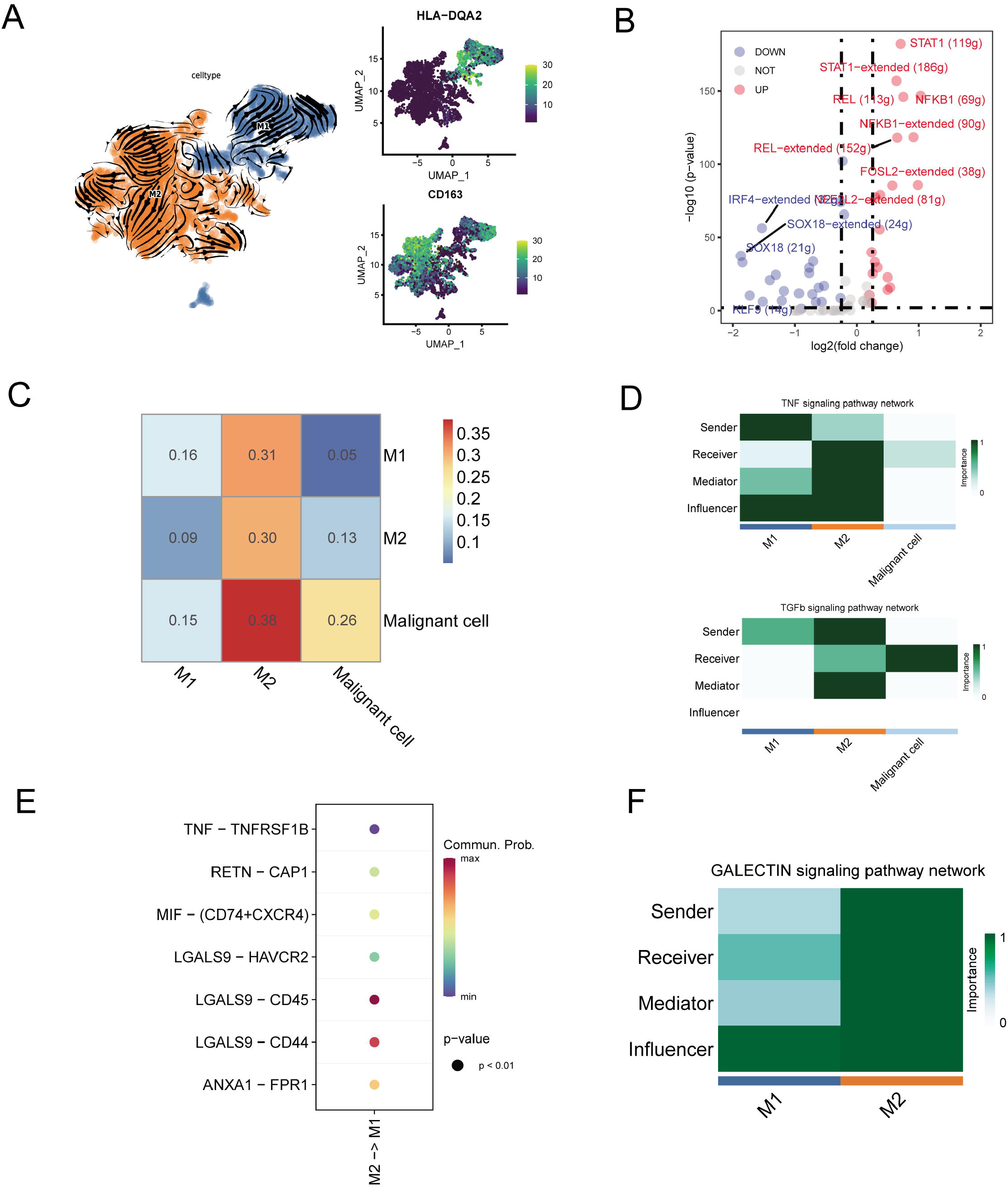
Macrophage subpopulations analysis. A. UMAP plot shows the RNA velocities and HLA-DQA2, CD163 expression patterns of macrophage subpopulations. A. The volcano map shows the difference regulators of macrophage subpopulations. C. Heatmap of the cell–cell interaction scores analyzed by CellChat. D. The role of each macrophage subpopulations cell in TNF and TGFb signaling pathway. E. Bubble chart showing ligand-receptor pairs secreted by M2 cells to M1 cells. F. The role of each macrophage subpopulations cell in GALECTIN signaling pathway.

### Lymphocyte subcluster analysis

B cell subpopulations are divided into Naïve B (IGHM+), memory B (CD27+), germinal center B (BCL6) and plasma cells (CD38+) (Figure5.A, FigureS6.A). The germinal center B cells and plasma cells are at the beginning and the end of the pseudo time respectively (Figure S6.B-C). We have discovered the molecular regulation relationship between various types of B cells and malignant cells (Figure5.B-C, FigureS6.D). Germinal center B cells can act on the CD44 receptors of malignant cells through LGALS9 ligand, while malignant cells act on the CD74+CXCR4, CD74+CD44 receptors of various types of B cells mainly through MIF ligand molecules.

**Figure 5.**
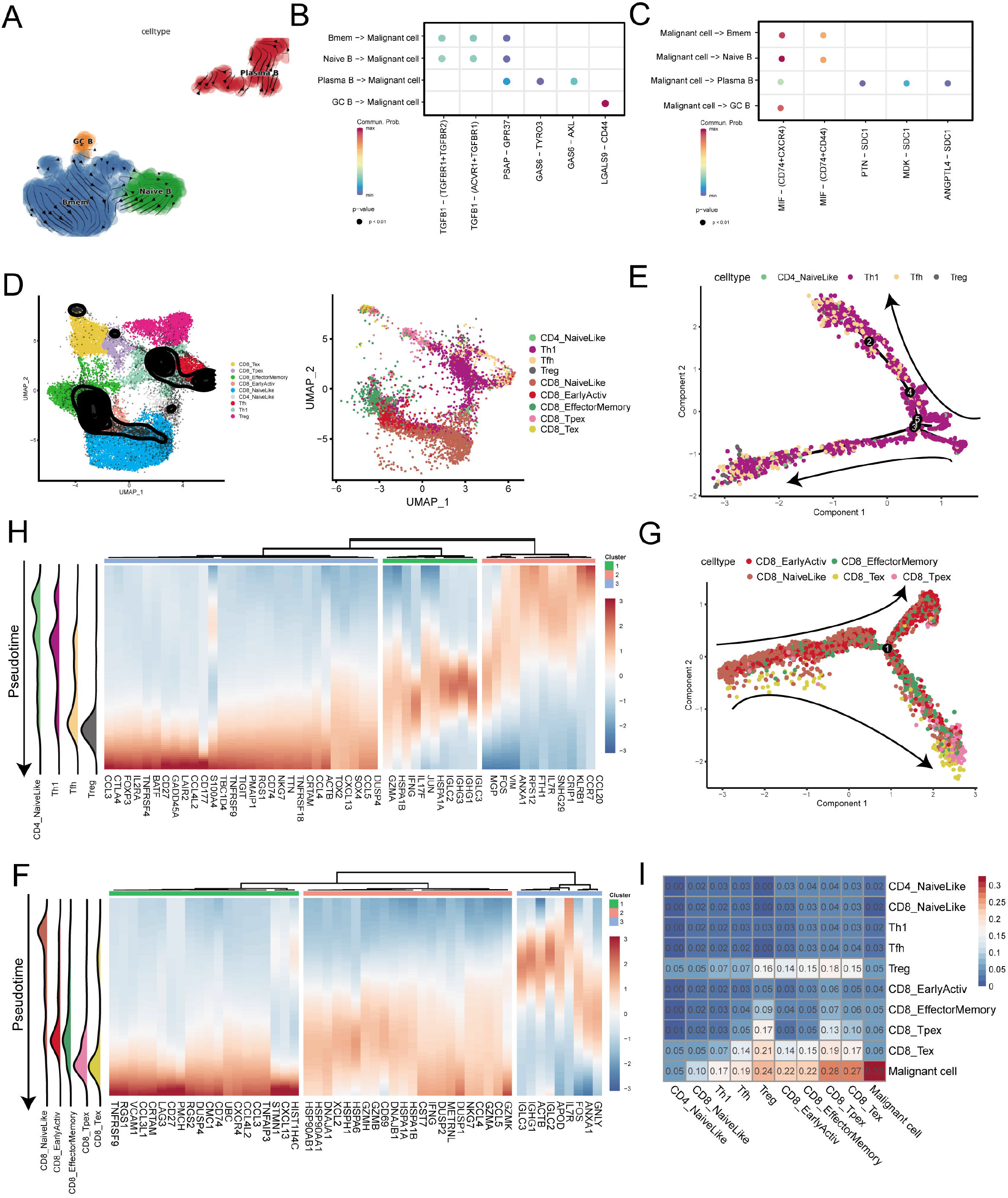
Lymphocyte cell subpopulations analysis. A. UMAP plot shows the RNA velocities and the latent time of B cell subpopulations. B. Bubble chart showing ligand-receptor pairs secreted by B cells to malignant cells. C. Bubble chart showing ligand-receptor pairs secreted by malignant cells to B cells. D. UMAP map shows the results of T cell subpopulation cell annotation by ProjecTILs. E. The arrangement of different CD4+ T subpopulations cells on pseudo-time trajectory. F. Heatmap shows dynamic gene expression patterns accompanying the differentiation of CD8+ T cells. G. The arrangement of different CD8+ T subpopulations cells on pseudo-time trajectory. H. Heatmap shows dynamic gene expression patterns accompanying the differentiation of CD8+ T cells. I. Heatmap of interaction strength between T cells and malignant cells.

Use ProjecTILs to annotate T cells in the data into 9 subpopulations, including CD4+ T cells (CD4_NaiveLike, Th1, Tfh, Treg) and CD8+ T cells (CD8_NaiveLike, CD8_EarlyActiv, CD8_EffectorMemory, CD8_Tpex, CD8_Tex) (Figure5.D). CD4+ T cells are gradually differentiated into Treg, which corresponds to the characteristic gene set (cluster3), represented by FOXP3 (Figure5.E-H). FOXP3 is a key regulator of regulatory T (Treg) cell gene expression, which can activate the expression of TNFRSF18, IL2RA and CTLA4, and inhibit the expression of IL2 and IFNG in association with the TF RUNX1 (15). The pseudotime trajectory depicts the difference in gene expression of CD8+ T cells from CD8_NaiveLike to exhausted (or prexhausted) T cells (CD8_Tpex, CD8_Tex) (Figure5.G). In the CD8+T pseudotime gene clustering heatmap, with the direction of pseudotime, CD8+ T cells gradually transformed from cluster3 genes with high expression of GNLY and IGHG1 to cluster2 genes with high expression of GZMK and IFNG. At the terminal cluster1, the exhausted genes represented by LAG3 is highly expressed (Figure5.F). Treg, CD8_Tpex and CD8_Tex interact strongly with malignant cells (Figure5.I).

### CTGF+ CAFs play a key role in TME

We found that CAFs accounted for the largest proportion of cells after malignant cells, and there were many interactions with malignant cells, which could play a key role in TME. CAFs were then subdivided into three subpopulations. In addition to the traditional iCAFs (RGS5+) and mCAFs (PDGFRA+) cell clusters, we also found new CTGF+ CAFs clusters (Figure6.A). Through GO enrichment of the differential genes of the three subpopulations, the cell functions of each cluster were found Figure6.B). iCAFs cluster was associated with BMP signaling pathway, extracellular matrix organization, extracellular structure organization etc. mCAFs cluster was associated with functions of muscle cell and tissue. CTGF+ CAFs cluster was associated with functions such as cell growth, response to hypoxia, protein folding, etc. SCENIC results showed that the TWIST1 motif was highly activated in iCAFs, and the TBX2 motif was highly activated in iCAFs. CTGF+ CAFs specifically activate NFATC4 and SOX10 motifs. (Figure6.C-D). With the development of tumors, the expression of CTGF+ CAFs gradually increase, which suggests that the secreted protein CTGF plays an important regulatory role in the evolution of malignant cells (Figure6.E-G).

**Figure 6.**
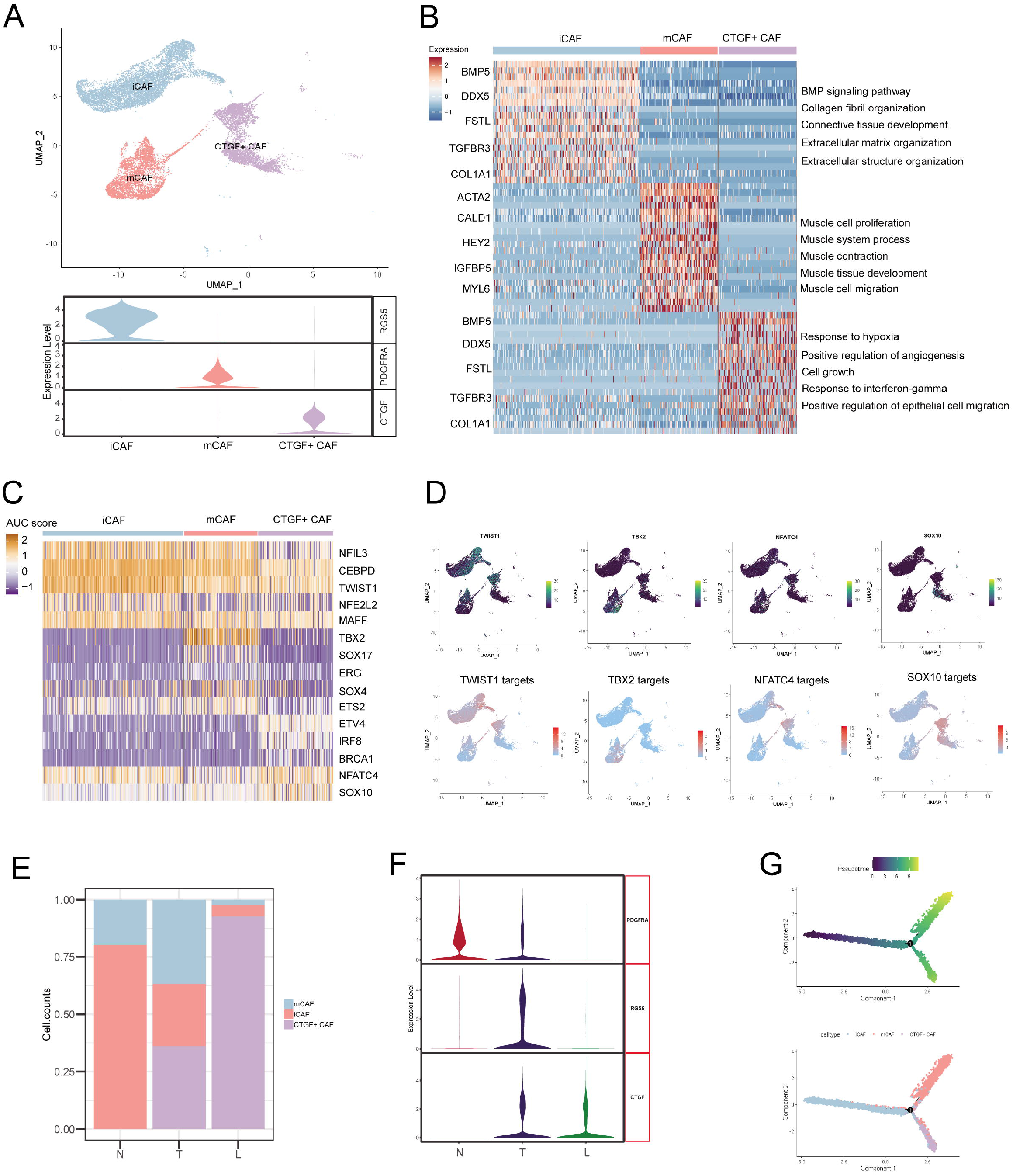
Fibrocyte subpopulations analysis. A. UMAP plot of Fibrocyte subpopulations and vlnplot of Fibrocyte subpopulations marker genes. B. Heatmap shows the subpopulations differentially expressed genes and enrichment to GO terms. C. Heatmap of the AUC scores of TF motifs estimated by SCENIC for each cell in Fibrocyte subpopulations. D. UMAP plot of TFs include TWIST1, TBX2, NFATC4, SOX10, and TFs targets expression level in each Fibrocyte. E. Distribution of fibrocyte subpopulations in normal, tumor, and lymph node metastasis samples. F. Fibroblast subpopulation markers expression level in normal, tumor, and lymph node metastasis samples. G. Developmental pseudo-time of fibrocyte and the arrangement of different subpopulation cells on pseudo-time trajectory.

## Discussion

In recent years, immunotherapy has become the most popular research field in medical tumor research, and to solve the problem that the curative effect of immunotherapy differs due to the difference of race and tumor type has become the frontier problem to be solved in this field. Unlike the epidemiological characteristics of AM in the western population, which is less than 10% in malignant melanoma patients, AM in the Asian population is as high as 70%, with poor prognosis and immune efficacy. Single-cell sequencing technology can more intuitively feel the composition and proportion of various types of cells in the microenvironment, and can find specific subgroups that cannot be paid attention to by traditional bulk sequencing. Tumor-infiltrated immune cells, including lymphocyte, TAM, myeloid-derived suppressor cell (MDSC), are important components of TME. On the one hand, these immune cells can kill tumor cells (such as CD8+ T cells and NK cells), and on the other hand, they can promote tumor development (16). Existing studies have shown that immune cells in TME can play a role in promoting tumor development through immunosuppression (17, 18), promoting angiogenesis (19), inhibiting apoptosis (20), secreting growth factors (21), helping tumors escape growth suppressor factors (22), promoting tumor metastasis (23), and changing energy metabolism (24). Taking macrophages as an example, TAM has two forms of macrophages, M1-type and M2-type. M1-type macrophages can inhibit and phagocytose tumor cells, while M2-type macrophages play immunosuppressive and tumor promoting roles. In addition to immune cell components, stromal components such as endothelial cells and fibroblasts are also important components of TME. By secreting transforming growth factor (TGF-β) (25) and vascular endothelial growth factor (VEGF) (26), tumor cells can induce and activate CAFs and endothelial cells, change tumor cell phenotype, reshape extracellular matrix (ECM), help generate blood vessels and lymphatic vessels, and accelerate the outward escape of tumor (27). In recent years, we have found that the stromal components in TME also affect the anti-tumor immune effect, which indicates that it is also necessary to target the related stromal cells in the process of tumor immunotherapy (28). Fibroblasts normally maintain the structure of tissue. However, in the early stage of tumor formation, many chemokines (IL-6, IL8, etc.) secreted by tumor cells can transform normal fibroblasts around the tumor into CAFs (29), gradually forming a microenvironment suitable for malignant proliferation and metastasis of tumor cells.

In view of the important role of TME in tumor progression, it has become an important therapeutic target. Taking the elimination of immunosuppression of CD8+T cells in TME as an example (30), The use of antibodies to block cytotoxic T lymphocyte antigen-4 (CTLA-4) and programmed cell death-1 (PD-1) immunodrugs have been marketed, and have achieved remarkable results in the treatment of melanoma (31), lymphoma (32), Mercer-cell carcinoma (33) and other tumors. Targeted drugs that block the activation of VEGF signaling pathway and inhibit tumor metastasis, such as Sorafenib and bevacizumab, have also been widely developed. However, although it has improved survival in patients with advanced stage, the efficacy is still limited and the response rate is low for the AMs that is more common in the Asian population. Therefore, a better understanding of the TME of AM could accelerate the discovery of new targets or combined therapeutic strategies, which in turn has important clinicopathological significance in helping clinicians select appropriate treatment regimens and predict outcomes. We generated a single-cell transcriptome landscape, uncovered the components of the microenvironment within AM tissues, and analyzed cell interactions.

In this study, we identified 9 cell types in AM microenvironment, among which malignant cells accounted for the largest proportion, followed by CAFs. Through cell communication analysis, we found that with the development of tumor, the interaction between malignant cells and macrophages, T cells, B cells and CAFs became more and more stronger. We suggest that malignant cells with high expression of TWIST1 have the highest degree of malignancy. We found that activation of STAT1, REL and NFKB1 mods led to this M2 polarization process. The results of cell interaction showed that there was strong cell interaction between M2 subpopulation and malignant cells. Bmen cells accounted for more in AM TME. FOXP3 was highly expressed at the end of CD4+ T cell evolution. And, LAG3 was highly expressed at the end of CD8+ T cell evolution.

In addition, CAFs are important in TME. Previous studies have found that CAFs regulate the angiogenesis by produce angiogenic factors, such as FGF2 and VEGFA, thereby providing essential for highly proliferating tumor cells. CAFs can also help tumor cells overcome immune surveillance by recruiting immunosuppressive cells, such as M2 and myeloid-derived suppressor cells (MDSC) (34, 35). In this AM data set, we have annotated a new subpopulation of fibroblasts, CTGF+ CAFs, which is a key factor in AM tumor progression. Using the area under the TF motif (AUC) score of each cell estimated by SCENIC, it is found that NFATC4 and SOX10 motif is specifically activated in CTGF+CAFs.

In summary, we determined the expression profile of cell subpopulations in AM and confirmed the characteristics of these tumor-related subpopulations. This cell atlas provides an in-depth understanding of cancer immunology and is an important resource for future drug discovery.

## Methods

### Patients and samples

All samples were obtained from the General Hospital of People’s Liberation Army, Beijing, China. Five primary AM tissues, two adjacent paracancerous tissues and a metastatic lymph node sample, were involved in this cohort. All experimental procedures were approved by the General Hospital of People’s Liberation Army, Beijing, China.

### Single-cell suspension preparation

Primary AM tissues, adjacent paracancerous tissues and a metastatic lymph node tissue were processed immediately after being obtained from AM patients. Single-cell suspensions with high cell viability (> 90%) were prepared using an automatic mild tissue processor (Miltenyi gentleMACS Dissociator). 1mg/ml collagenase and 1mg/ mL elastase were prepared in a ratio of 1:4 and preheated at 37°C. Every sample was cut into small pieces (<1□mm in diameter). The tissues and digestive juices are put into the C tube or M tube matching the instrument, and the corresponding tube cover is covered. Then the tube cover is inverted and installed on the disintegrator. Select the gentleMACS program from the menu, set the steering and speed, and press the Start button to start. After operation, remove the tube, open the tube cover and take out the cell suspension. The cells were washed twice with buffer solution and then resuspended to 800-1200 cells /ul. The cells were counted by Trypan blue or fluorescent reagents and corresponding counting instruments. Cell suspensions with cell viability ≥90% and aggregation rate less than or equal to 5% could be used for sequencing.

### Droplet-based single-cell sequencing

Chromium Single Cell 3’ Library and Gel Bead Kit V3 (10× Genomics, 1000075) were used to prepare barcoded scRNA-seq libraries according to the manufacturer’s protocol. Single-cell suspensions were loaded onto a Chromium Single-Cell Controller Instrument (10× Genomics), in order to generate single-cell gel beads in emulsions (GEMs) according to the manufacturer’s protocol. Approximately 8,000 cells were added to each channel, to capture 5000 cells per library. Captured cells were lysed and the released RNA were barcoded through reverse transcription in individual GEMs. Using a S1000TM Touch Thermal Cycler (Bio Rad) to reverse transcribe, the GEMs were programed at 53°C for 45 min, followed 85°C for 5 min, and hold at 4°C. The cDNA was generated and then amplified, and the quality was assessed using the Agilent 4200. Every library was sequenced on a Illumina Novaseq 6000 sequencer with a sequencing depth of at least 100,000 reads per cell and 150 bp (PE150) paired-end reads were generated (performed by CapitalBio, Beijing).

### Raw data processing and quality control

Cell Ranger (version 3.3.0) was used to process the raw data, demultiplex cellular barcodes, map reads to the transcriptome, and down-sample reads (as required to generate normalized aggregate data across samples). Raw gene expression matrices with unique molecular identifier (UMI) generated by Cell Ranger were imported into Seurat (v.4.0.0) (36). Cells with percentage of mitochondria reads ≥ 25%, unique genes ≤ 500 were considered low-quality cells and were removed. DoubletFinder was used to eliminate potential doublets. Finally, 61726 single cells remained, and they were applied in downstream analyses (Figure S1.A-B). After removed potential batch effect, 20 principal components (PC) were used to principal component analysis (PCA), and UMAP was used tononlinear dimension reduction and visualization.

### Cell type annotation

The CopyKAT (10) package was used to detect the CNVs in cells and to recognize real cancer cells with default parameters. Aneuploid cells were defined as malignant cells, and diploid cells were annotated by SingleR and classified according to annotation results (11).

### Trajetory and RNA velocity analysis

RNA velocity and pseudotime analysis were performed with Monocle2 (37) and scVelo (37). ScVelo is a Python(v3.9.0)-based computational analysis tool. Other data analyses were completed in R (v4.0.0) environment.

### Simultaneous gene regulatory network analysis

SCENIC is a new computational method used in the construction of regulatory networks and in the identification of different cell states from scRNA-seq data (13). To measure the difference between cell clusters based on TFs or their target genes, SCENIC was performed on all single cells, and the preferentially expressed regulons were calculated by the Limma package (38). Only regulons significantly upregulated or downregulated in at least one cluster, with adj. p-value□<□0.05, were involved in further analysis.

### Cell-cell communication analysis

Cell contact patterns were constructed by CellChat (v.0.0.2) (12). CellChat uses the gene expression data as input and combines the interactions of ligand receptors and their cofactors to simulate cell-to-cell communication. It can identify communication patterns, and predict the function of understudied pathways and the key signaling events between spatially co-localized cell populations.

### Functional enrichment analysis

Differentially expressed genes (DEGs) of cell subgroups were recognized by the findmarker function provided by Seurat. Log2|FC| > 0.5 and adj.p.val□<□0.01 were used as the cut-off criteria. ClusterProfiler was used for GO/KEGG analysis (Gene Ontology/ Kyoto Encyclopedia of Genes and Genomes) (39–41). GO was used to describe gene functions along three aspects: biological process (BP), cellular component (CC) and molecular function (MF). The KEGG was searched for pathways at the significance level set at p<0.05.

## Supplementary material

Figure S1 Data quality control

A. Bar plot of doublet and singlet number. B. Vlnplot of feature counts, UMI counts and percentage of mitochondrial genes in cells after quality control. C. UMAP plots of feature counts, UMI counts and percentage of mitochondrial genes in cells after quality control. D. UMAP plot display sample cell distribution. E. Bar plot show cell number of per sample after quality control.

Pie chart of the cell number percentage of the two batches. B. Correlation analysis of the two batches by the Spearman correlation coefficient of all detected genes.

Figure S2 Cell annotation

A. Heatmap plot showing the Copykat result. B. Pie chart of the diploid and aneuploid cell number percentage. C. Bar plot of doublet and singlet number. D. Heatmap plot showing the SingleR result. E. Chord diagram shows the correspondence between cell annotation results and cell cluster results. F. Cell type pearson correlation coefficient heatmap.

Figure S3 Cell communication

A. Bar graph show the conserved and context-specific signaling pathways. B. The role of each cell type in KIT and WNT signaling pathway of normal samples. C. The role of each cell type in NT, ncWNT, IL1 and GDF signaling pathway of tumor samples. D. The role of each cell type in CHEMERIN, NRG and PSAP signaling pathway of lymph node metastasis sample. E. Interaction strength between malignant cells and other types of cells.

Figure S4 Lymphocyte cell subpopulations gene expression patterns

A. UMAP plot of each B cell BCL6, IGHM, CD27 and CD38 expression level. B. The latent time of B cells. C. Heatmap shows dynamic gene expression patterns accompanying the evolution of B cells. D. Heatmap of interaction strength between B cells and malignant cells.

## Notes

### Competing Interest Statement

The authors have declared no competing interest.

## Reference

1. Swick JM, Maize JC, Sr. Molecular biology of melanoma. J Am Acad Dermatol. 2012;67(5):1049–54.

2. Wahid M, Jawed A, Mandal RK, Dar SA, Akhter N, Somvanshi P, et al. Recent developments and obstacles in the treatment of melanoma with BRAF and MEK inhibitors. Crit Rev Oncol Hematol. 2018;125:84–8.

3. McKean MA, Amaria RN. Multidisciplinary treatment strategies in high-risk resectable melanoma: Role of adjuvant and neoadjuvant therapy. Cancer Treat Rev. 2018;70:144–53.

4. Cormier JN, Xing Y, Ding MC, Lee JE, Mansfield PF, Gershenwald JE, et al. Ethnic differences among patients with cutaneous melanoma. Arch Intern Med. 2006;166(17):1907–14.

5. Chi ZH, Li SM, Sheng XN, Si L, Cui CL, Han M, et al. Clinical presentation, histology, and prognoses of malignant melanoma in ethnic Chinese: A study of 522 consecutive cases. Bmc Cancer. 2011;11.

6. Nakamura Y, Namikawa K, Yoshino K, Yoshikawa S, Uchi H, Goto K, et al. Anti-PD1 checkpoint inhibitor therapy in acral melanoma: a multicenter study of 193 Japanese patients. Ann Oncol. 2020;31(9):1198–206.

7. Rose AAN, Armstrong SM, Hogg D, Butler MO, Saibil SD, Arteaga DP, et al. Biologic subtypes of melanoma predict survival benefit of combination anti-PD1+anti-CTLA4 immune checkpoint inhibitors versus anti-PD1 monotherapy. J Immunother Cancer. 2021;9(1).

8. Wan LL, Pantel K, Kang YB. Tumor metastasis: moving new biological insights into the clinic. Nat Med. 2013;19(11):1450–64.

9. Tang FC, Barbacioru C, Wang YZ, Nordman E, Lee C, Xu NL, et al. mRNA-Seq whole-transcriptome analysis of a single cell. Nat Methods. 2009;6(5):377–U86.

10. Gao RL, Bai SS, Henderson YC, Lin YY, Schalck A, Yan Y, et al. Delineating copy number and clonal substructure in human tumors from single-cell transcriptomes. Nat Biotechnol. 2021;39(5):599–+.

11. Aran D, Looney AP, Liu LQ, Wu E, Fong V, Hsu A, et al. Reference-based analysis of lung single-cell sequencing reveals a transitional profibrotic macrophage. Nat Immunol. 2019;20(2):163–+.

12. Jin SQ, Guerrero-Juarez CF, Zhang LH, Chang I, Ramos R, Kuan CH, et al. Inference and analysis of cell-cell communication using CellChat. Nat Commun. 2021;12(1).

13. Aibar S, Gonzalez-Blas CB, Moerman T, Van AHT, Imrichova H, Hulselmans G, et al. SCENIC: single-cell regulatory network inference and clustering. Nat Methods. 2017;14(11):1083–+.

14. Donn RP, Ray DW. Macrophage migration inhibitory factor: molecular, cellular and genetic aspects of a key neuroendocrine molecule. J Endocrinol. 2004;182(1):1–9.

15. Wu YQ, Borde M, Heissmeyer V, Feuerer M, Lapan AD, Stroud JC, et al. FOXP3 controls regulatory T cell function through cooperation with NFAT. Cell. 2006; 126(2):375–87.

16. Hanahan D, Weinberg RA. Hallmarks of Cancer: The Next Generation. Cell. 2011;144(5):646–74.

17. Ruffell B, DeNardo DG, Affara NI, Coussens LM. Lymphocytes in cancer development: Polarization towards pro-tumor immunity. Cytokine Growth F R. 2010;21(1):3–10.

18. Joyce JA, Fearon DT. T cell exclusion, immune privilege, and the tumor microenvironment. Science. 2015;348(6230):74–80.

19. DeNardo DG, Johansson M, Coussens LM. Immune cells as mediators of solid tumor metastasis. Cancer Metast Rev. 2008;27(1):11–8.

20. Chen Q, Zhang XHF, Massague J. Macrophage Binding to Receptor VCAM-1 Transmits Survival Signals in Breast Cancer Cells that Invade the Lungs. Cancer Cell. 2011;20(4):538–49.

21. Balkwill F, Charles KA, Mantovani A. Smoldering and polarized inflammation in the initiation and promotion of malignant disease. Cancer Cell. 2005;7(3):211–7.

22. Lu PF, Takai K, Weaver VM, Werb Z. Extracellular Matrix Degradation and Remodeling in Development and Disease. Csh Perspect Biol. 2011;3(12).

23. Kessenbrock K, Plaks V, Werb Z. Matrix Metalloproteinases: Regulators of the Tumor Microenvironment. Cell. 2010;141(1):52–67.

24. Buck MD, Sowell RT, Kaech SM, Pearce EL. Metabolic Instruction of Immunity. Cell. 2017;169(4):570–86.

25. Noma K, Smalley KSM, Lioni M, Naomoto Y, Tanaka N, El-Deiry W, et al. The essential role of fibroblasts in esophageal squamous cell carcinoma-induced angiogenesis. Gastroenterology. 2008;134(7):1981–93.

26. Olofsson B, Jeltsch M, Eriksson U, Alitalo K. Current biology of VEGF-B and VEGF-C. Curr Opin Biotech. 1999;10(6):528–35.

27. Folkman J. What Is the Evidence That Tumors Are Angiogenesis Dependent. Jnci-J Natl Cancer I. 1990;82(1):4–6.

28. Turley SJ, Cremasco V, Astarita JL. Immunological hallmarks of stromal cells in the tumour microenvironment. Nat Rev Immunol. 2015;15(11):669–82.

29. Zhang Y, Liu ZY, Yang X, Lu WQ, Chen YL, Lin YB, et al. H3K27 acetylation activated-COL6A1 promotes osteosarcoma lung metastasis by repressing STAT1 and activating pulmonary cancer-associated fibroblasts. Theranostics. 2021;11(3):1473–92.

30. Topalian SL, Taube JM, Anders RA, Pardoll DM. Mechanism-driven biomarkers to guide immune checkpoint blockade in cancer therapy. Nat Rev Cancer. 2016;16(5):275–87.

31. Eroglu Z, Kim DW, Wang XY, Camacho LH, Chmielowski B, Seja E, et al. Long term survival with cytotoxic T lymphocyte-associated antigen 4 blockade using tremelimumab. Eur J Cancer. 2015;51(17):2689–97.

32. Robert C, Long GV, Brady B, Dutriaux C, Maio M, Mortier L, et al. Nivolumab in Previously Untreated Melanoma without BRAF Mutation. New Engl J Med. 2015;372(4):320–30.

33. Engels EA. Epidemiologic perspectives on immunosuppressed populations and the immunosurveillance and immunocontainment of cancer. Am J Transplant. 2019;19(12):3223–32.

34. Flavell RA, Sanjabi S, Wrzesinski SH, Licona-Limon P. The polarization of immune cells in the tumour environment by TGFbeta. Nat Rev Immunol. 2010;10(8):554–67.

35. Yang XG, Lin YL, Shi YH, Li BJ, Liu WR, Yin W, et al. FAP Promotes Immunosuppression by Cancer-Associated Fibroblasts in the Tumor Microenvironment via STAT3-CCL2 Signaling. Cancer Res. 2016;76(14):4124–35.

36. Stuart T, Butler A, Hoffman P, Hafemeister C, Papalexi E, Mauck WM, et al. Comprehensive Integration of Single-Cell Data. Cell. 2019;177(7):1888–+.

37. Qiu XJ, Mao Q, Tang Y, Wang L, Chawla R, Pliner HA, et al. Reversed graph embedding resolves complex single-cell trajectories. Nat Methods. 2017;14(10):979–+.

38. Ritchie ME, Phipson B, Wu D, Hu YF, Law CW, Shi W, et al. limma powers differential expression analyses for RNA-sequencing and microarray studies. Nucleic Acids Res. 2015;43(7).

39. Ashburner M, Ball C, Blake J, Botstein D, Butler H, Cherry J, et al. Gene ontology: tool for the unification of biology. The Gene Ontology Consortium. Nature genetics. 2000;25(1):25–9.

40. Kanehisa M, Goto S, Furumichi M, Tanabe M, Hirakawa M. KEGG for representation and analysis of molecular networks involving diseases and drugs. Nucleic acids research. 2010;38:D355–60.

41. Yu G, Wang L, Han Y, He Q. clusterProfiler: an R package for comparing biological themes among gene clusters. Omics : a journal of integrative biology. 2012;16(5):284–7.

