## Supplementary figures and images for "Mapping of single-cell landscape of acral melanoma and analysis of molecular regulatory network of tumor microenvironment"

### Supplemental Figure 1

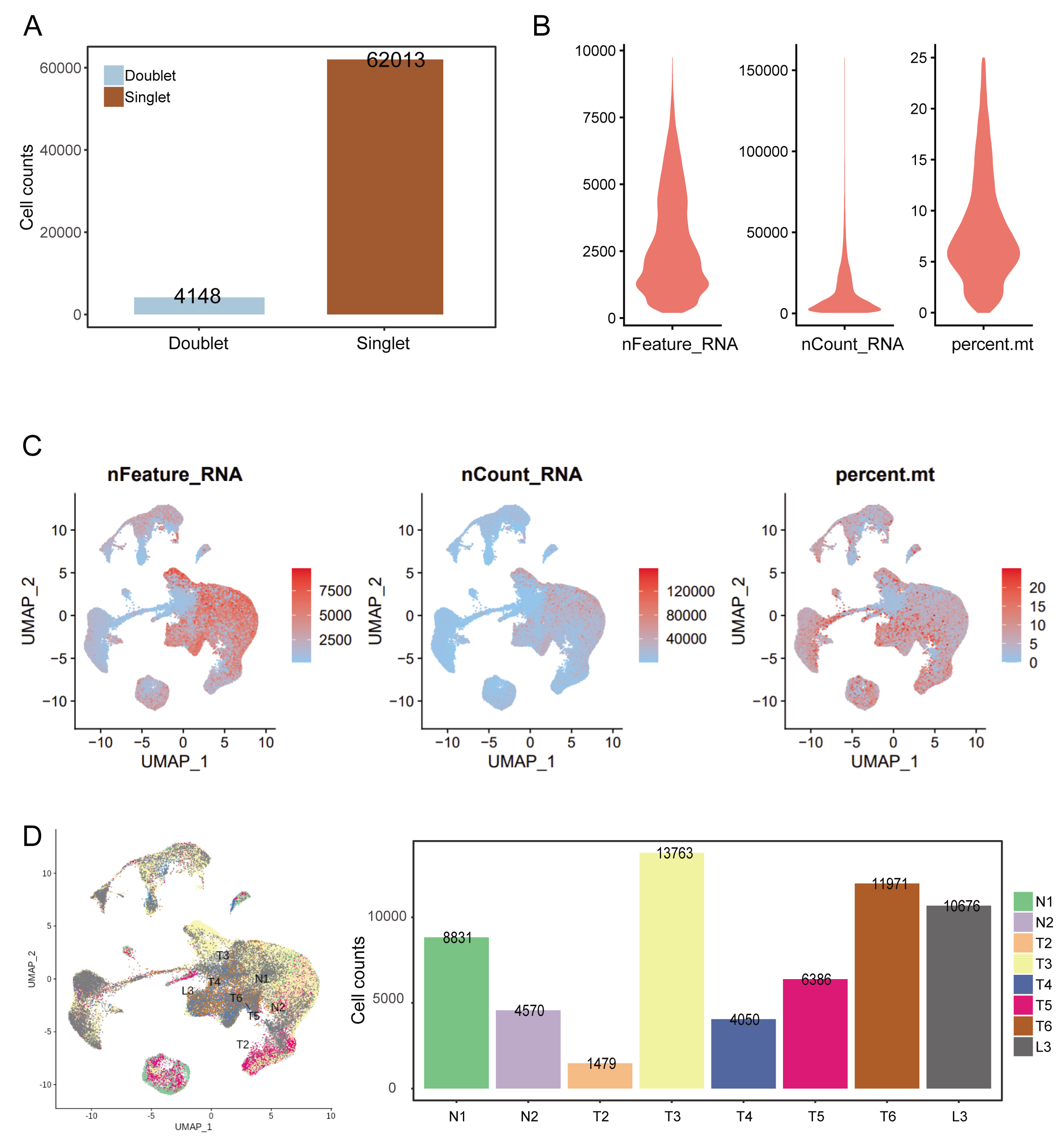

### Supplemental Figure 2

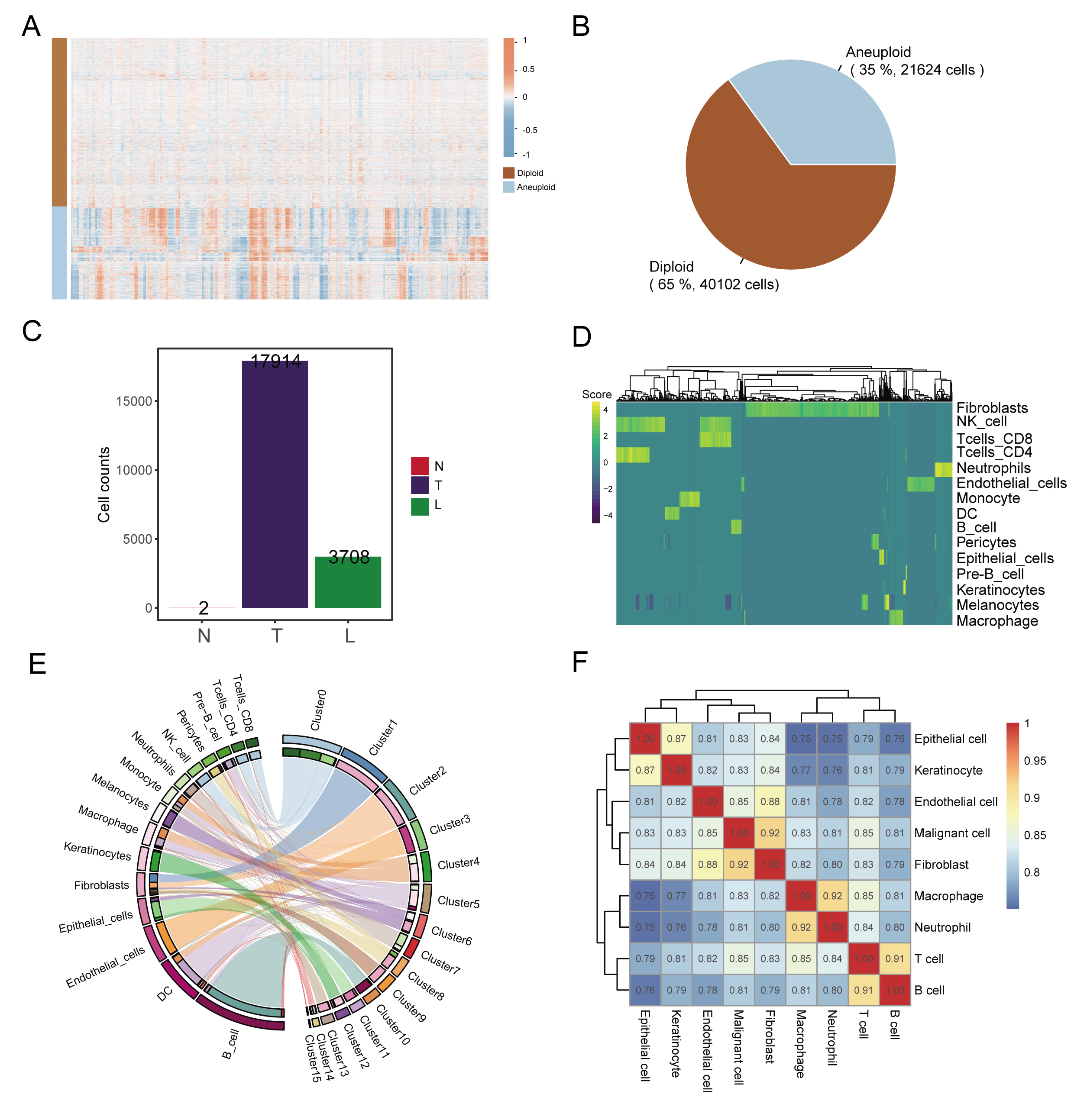

### Supplemental Figure 3

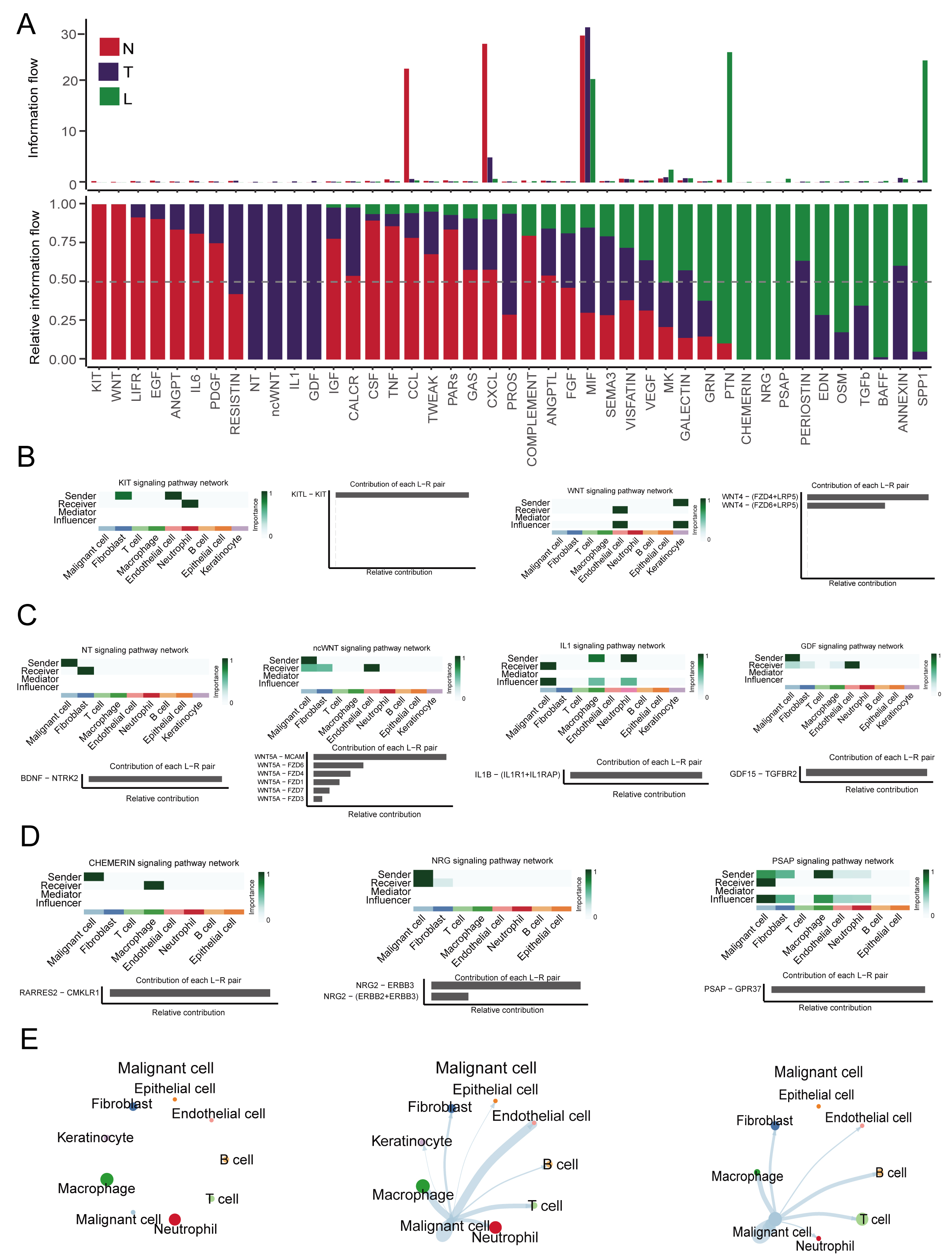

### Supplemental Figure 4

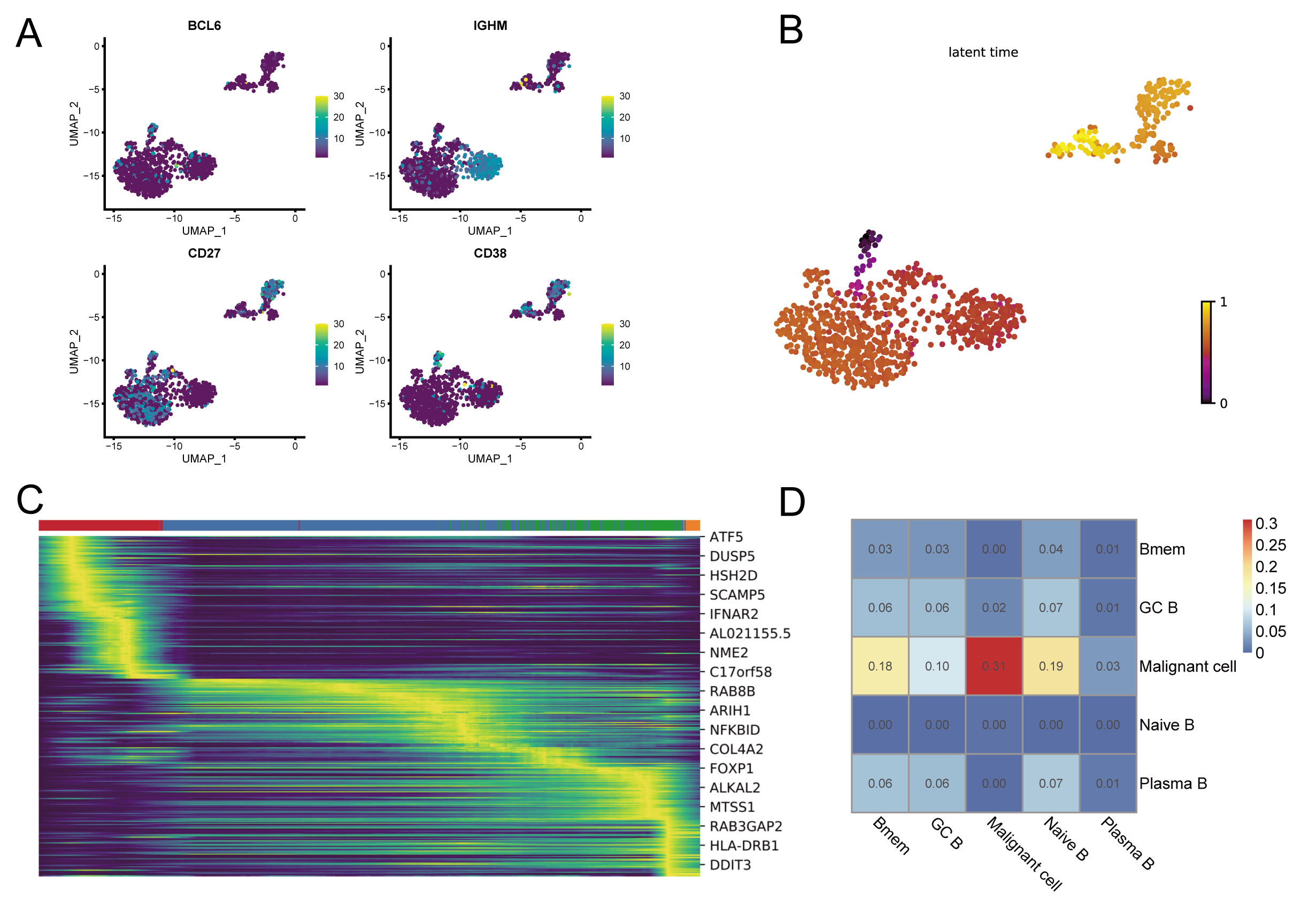
